# Characterising *Staphylococcus aureus* genomic epidemiology with Multilevel Genome Typing

**DOI:** 10.1101/2024.05.09.593273

**Authors:** Liam Cheney, Michael Payne, Sandeep Kaur, Genevieve McKew, Ruiting Lan

## Abstract

*Staphylococcus aureus* is a major source of both hospital and community acquired infections, and is the leading source of skin and soft tissue infections worldwide. Advances in whole genome sequencing (WGS) technologies have recently generated large volumes of *S. aureus* WGS data. The timely classification of *S. aureus* WGS data with genomic typing technologies has the potential to describe detailed genomic epidemiology at large and small scales. In this study, a multilevel genome typing (MGT) scheme comprised of 8 levels of multilocus sequence typing schemes of increasing resolution was developed for *S. aureus* and used to analyse 50,481 publicly available genomes. Application of MGT to *S. aureus* epidemiology was showcased in three case studies. Firstly, the population structure of the globally disseminated sequence type ST8 were described by MGT2, which was compared with *Spa* typing. Secondly, MGT was used to characterise MLST ST8 - PFGE USA300 isolates that colonised multiple body sites of the same patient. Unique STs from multiple MGT levels were able to group isolates of the same patient, and the highest resolution MGT8 separated isolates within a patient that varied in predicted antimicrobial resistance. Lastly, MGT was used to describe the transmission of MLST ST239 - SCC*mec* III throughout a single hospital. MGT STs were able to describe both isolates that had spread between wards and also isolates that had colonised different reservoirs within a ward. The *S. aureus* MGT describes large- and small-scale *S. aureus* genomic epidemiology with scalable resolutions using stable and standardised ST assignments. The *S. aureus* MGT database is online (https://mgtdb.unsw.edu.au/staphylococcus) and is capable of tracking new and existing clones to facilitate the design of new strategies to reduce the global burden of *S. aureus* related diseases.

## Introduction

*Staphylococcus aureus* causes infections in both hospitalised individuals and those without clinical associations who are otherwise considered “healthy” (1). In 2017, *S. aureus* caused over 120,000 bloodstream infections and was associated with 20,000 deaths in the USA alone A hallmark of *S. aureus* is its ability to acquire new mechanisms of antimicrobial resistance (AMR). The first report of methicillin resistant *S. aureus* (MRSA) was in 1961 which was followed by a series of epidemic waves that have each acquired additional AMR The spread of these epidemic waves was predominantly reported in individuals associated with hospitals, and isolates were referred to as hospital associated-MRSA (HA-MRSA) (4). In the 1990s, there were reports of individuals not associated with clinical settings carrying MRSA. These cases were defined as community associated MRSA (CA-MRSA). Recent epidemiological surveillance has reported the spread of CA-MRSA throughout hospital settings, and CA-MRSA has been predicted to displace existing HA-MRSA in most countries as the leading cause of MRSA (5, 6).

Early applications of multilocus sequence typing (MLST) have retrospectively studied the global spread of HA-MRSA. One such example is ST239 that between 1970 and 1990 spread globally to cause multiple epidemics of HA-MRSA (7). MLST has also characterised the emergence and spread of CA-MRSA. Comparative study between HA-MRSA and CA-MRSA have shown that CA-MRSA are more diverse in sequence types (STs) and geographically restricted (8).

*S. aureus* WGS data is most often analysed through phylogenomic reconstruction. The comparison of geographically diverse isolates sampled over long periods has uncovered the origins of globally disseminated clones (9–12). For example, the study of ST8 sampled across the globe identified its European origin and transmission into North America in the 2000s (9). Once ST8 was in North America, phylogenetic analyses confirmed subsequent parallel epidemics of ST8-USA300 and ST8-USA400 CA-MRSA in North and South America, respectively (13).

While investigating *S. aureus* genomic epidemiology using phylogenetic approaches has provided high resolution comparisons, phylogenetics without the further implementation of genotyping algorithms cannot classify isolates into “types”. The classification of *S. aureus* has the potential to be revolutionised by the development of WGS-based typing technologies. In 2014, a *S. aureus* cgMLST scheme was developed that offered high resolution typing and a standardised nomenclature (14). The cgMLST has been sporadically used in smaller-scale studies such as outbreaks in neonatal wards and household transmission (15–17). cgMLST for each of these investigations has distinguished epidemiologically indistinguishable isolates and allowed the clustering of cgSTs to describe population structure. However, cgMLST for the species-wide classification is hampered by a lack of flexibility in typing resolution. The existing cgMLST applications required setting allele thresholds to cluster isolates. Selecting different allele thresholds between investigations would prevent establishing a standardised nomenclature, and to date, no such allele thresholds have been rigorously investigated and defined. Furthermore, the ability to vary the typing resolution is essential to describing large-scale epidemiology, such as global dissemination of ST8, and small-scale epidemiology, such as ST8-USA300 outbreaks within a hospital setting (9, 16).

We previously developed a new method called multilevel genome typing (MGT) which has been applied to several organisms (18–21). MGT is comprised of a series of MLST schemes of different sizes with scalable resolution. This study developed a *S. aureus* MGT that (1) offered flexibility in typing resolution to describe large and small scale genomic epidemiology and (2) established a standardised nomenclature for the clear communication of *S. aureus* types between investigations.

## Methods

### Dataset curation

Paired-end short read datasets for 66,238 genomes that were sequenced using any of the Illumina Genome Analyser, HiSeq, MiSeq, Next Seq and NovaSeq platforms were downloaded on the 15^th^ of July 2019 from the Sequence Read Archive of the National Centre for Biotechnology Information (NBCI). Additionally, publicly available ‘year of isolation’ metadata, where available, was downloaded from NCBI BioSample. Read sets were screened for contamination with Kraken (v1.0.0) and read sets with more than 20% non-*S. aureus* reads were removed (22). Assemblies were generated with the MGTdb pipeline which trimmed reads with Trimmomatic (v0.39.0), performed reference based assembly with Shovill (v1.0.9) and Skesa (v.2.3.0), and calculated assembly quality metrics with Quast (v5.2.0) (23–26). Assembles were quality filtered using the thresholds in Table 1. The quality filtered species dataset had 50,481 genomes (Supplementary Table 1).

**Table 1.** Quality filtering metrics for *S. aureus* assemblies.

| Filtering Metric | Range |
| --- | --- |
| Genome Size (mb) | Between 2.38 and 3.22 |
| N50 | > 50,000 |
| Average Contig Size (base pairs) | > 15,000 |
| Number of Contigs | < 150 |
| GC (%) | Between 32.5 and 33 |

### MGT Design

MGT1 to MGT8 loci were selected from a previously defined *S. aureus* core genome (14). MGT2-7 loci were selected based on previously published methods (20, 27, 28). The MGT design methodology included four stages: firstly, calculating the size of each level, secondly, calculating selection criteria for each core locus, and thirdly, separating the core loci into preferences, and fourthly, selecting core loci to fill the levels (Supplementary Methods, Supplementary Datasets, Supplementary Scripts).

MGT typing of the *S. aureus* species dataset.

The species dataset (*n=50,481*) was processed by the MGT pipeline (28). The reference-alleles of core loci were selected using the available complete reference genome *S. aureus* subsp. *aureus* COL (NCBI SRA accession: GCF_000012045). The allele calling pipeline of the MGT used the following thresholds: a maximum of 16 SNPs within a 40-base sliding window, 80% BLAST nucleotide identity, and 80% BLAST high-scoring segment pair. The MGT used the 16 SNP and 40-base sliding window to handle misaligned regions of DNA that can occur in the MGT allele calling pipeline.

### Allele-based phylogenetic construction

Allele-based phylogenies were constructed for the isolates of MGT1 ST8. Allele profiles from MGT8 (equivalent to cgMLST) were used to generate a phylogeny GrapeTree (v1.0.0), using the rapid neighbour joining algorithm (29). The MLST ST8 phylogeny was compared to MGT classification of MLST ST8. The isolates of the phylogeny generated by GrapeTree were coloured by their assigned MGT2 ST.

### SNP-based phylogenetic construction

Two SNP-based phylogenies were generated, one each for populations of ST239-SCC*mec* III and ST8-USA300, respectively (30, 31). Except for the chosen reference genome, generating SNP-based phylogenies for each of these populations used the same process. The reference genome *S. aureus* JKD6008 (NCBI SRA accession: GCA_000145595) was used to generate ST239-SCC*mec* III phylogeny, and the reference genome *S. aureus* TCH1516 (NCBI SRA accession: GCF_000017085) was used to ST8-USA300 phylogeny. To generate both phylogenies, SNPs were called against the reference genome and a SNP alignment was generated using the default Snippy-Core settings (v4.6.0) (32). SNPs under the influence of recombination were predicted and removed from the SNP alignment with RecDetect (v6.1) (33). RecDetect predicted recombination SNPs using a strict recombination prediction model. IqTree (v2.0.4) processed the SNP alignment to create a maximum likelihood phylogeny using 1,000 bootstraps and automatic selection of model parameters (34). Additional metadata was visualised on the phylogeny using iTOL (v6) (35).

### *In silico Spa* and SCCmec typing

All *spa* type comparisons used *spa* types that were predicted *in silico* by SpaTyper (v0.3.3) with default settings (36). SCCmec types were predicted using <u>staphopia-sccmec</u> (v1.0.0) with default settings (37). Note that that the publicly available version of Staphopia used in this study can only identify SCCmec types I-VII and recently described types (VIII-XIV) were not identifiable (38).

## Results

### The *S. aureus* MGT consists of eight levels

The *S. aureus* MGT consisted of eight levels with increasing numbers of loci (**Table 2**). MGT1 was the traditional seven-gene *S. aureus* MLST scheme and MGT8 was the species cgMLST which had 1,724 core loci. The intermediate MGT levels MGT2, 3, 4, 5, 6 and 7 each had 21, 38, 74, 187, 369 and 747 loci, respectively (**Table 2**). Of the global dataset (*n=50,481*), 98.73% (49,838/50,481) of the isolates were assigned an ST at all MGT levels. The remaining 1.27% (643/50,481) of isolates were not assigned an ST at one of more of the eight MGT levels.

**Table 2.** Overview of the *S. aureus* MGT levels.

| MGT Level | Number of Loci | No. Major STs* | Isolates in Major STs | percentage of isolates in Major STs | Average year span of Major STs | % of isolates in continent specific major STs^ |
| --- | --- | --- | --- | --- | --- | --- |
| MGT1 | 7 | 144 | 46621 | 96.14 | 17.29 | 30.22 |
| MGT2 | 21 | 338 | 33983 | 70.95 | 11.36 | 50.22 |
| MGT3 | 38 | 450 | 26234 | 54.78 | 8.6 | 84.27 |
| MGT4 | 74 | 429 | 17788 | 37.15 | 6.84 | 92.98 |
| MGT5 | 187 | 339 | 9721 | 20.30 | 4.07 | 96.48 |
| MGT6 | 369 | 243 | 6265 | 13.08 | 1.75 | 96.17 |
| MGT7 | 747 | 173 | 4095 | 8.55 | 1.07 | 99.4 |
| MGT8 | 1,723 | 113 | 2436 | 5.08 | 0.3 | 100 |

\*Major STs = STs with more than 10 isolates assigned to them
^ Continent specific STs = > 80% of assigned isolates from one continent

### Epidemiology of the *S. aureus* species described using MGT

The division of the *S. aureus* population structure by MGT was examined using the species dataset (50,481 isolates, Supplementary table 1). Of this dataset 26,416 (52.3%) had year metadata, 27,046 (53.6%) had country metadata and 25,596 (50.7%) had both. The average number of years that major STs were sampled for showed a clear trend of shorter timespan in higher resolution MGT levels (Table 1, Supp figure 1) demonstrating the potential epidemiological usefulness of each level in describing short and long lived clones.

**Figure 1.**
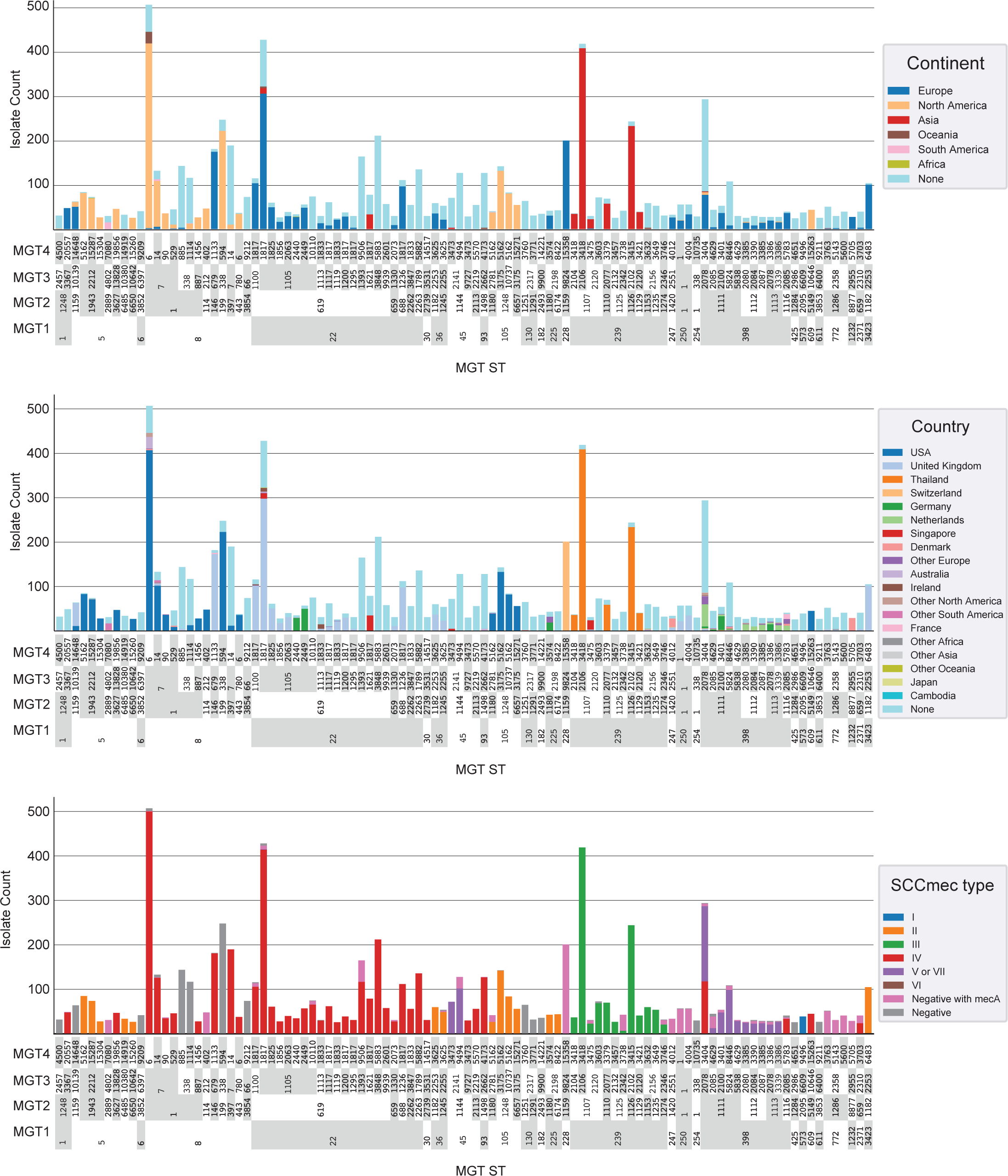
Geographical and MRSA diversity of the *S. aureus* species described using the 100 largest MGT4 STs. For all plots each column is one MGT4 ST. Y axis is isolate count in each ST. MGT1, MGT2 and MGT3 STs are shown below each MGT4 ST and grouped to allow examination of STs at these levels. A. The proportion of each continent in each MGT4 ST is represented by column colours. B. The proportion of each country in each MGT4 ST is represented by column colours. Countries with fewer than 200 isolates were grouped into “Other Continent” categories for clarity. C. SCCmec types assigned in each MGT4 ST.

To determine which level would best describe geographical trends we identified major STs (STs with more than 10 isolates assigned to them) with over 80% of their isolates from one continent. We then identified the lowest resolution level where more than 90% of isolates were within these continent-specific STs (Table 1). The value for MGT4 was 92.98%, therefore this level was selected to examine the distribution of continent-specific STs. The 100 largest MGT4 STs contained 7952 isolates, 4343 of which included continent metadata (Figure 1A). These STs show distinct distributions in both continent and country (Figure 1B). Of the largest 100 STs 26 contained no continent metadata and the remaining 74 were continent specific. Of these, 63 were specific to one country while 11 were found in more than one country in the same continent.

SCCmec types were also assigned to all isolates and their distribution among the top 100 MGT4 STs are shown (Figure 1C). In many cases all isolates within an MGT1 STs were MRSA (or at least contained mecA) and were a single SCCmec type (e.g. MGT1 ST22 and SCCmec IV). In other cases MGT2 or MGT3 STs are required to describe groups of isolates that are either exclusively MSSA or are MRSA and contain a single SCCmec type (e.g. MGT3 ST7: type IV, MGT3 ST338:MSSA and MGT3 ST2141: type ‘V or VII’). At MGT4 only ST 3404 contains two SCCmec types (IV and ‘V or VII’).

### Using MGT to describe MGT1 ST8 isolates

We selected MGT1 ST8 (traditional MLST ST8) to demonstrate the application of MGT to *S. aureus* epidemiology. Of the 50,481 isolates typed, 4,388 isolates were MGT1 ST8 and had associated collection year metadata. As MGT levels increased in resolution the number of STs at each level increased while the size of STs decreased (Figure 2). The largest MGT2 ST (ST1) was assigned to 37.58% (1,649/4,388) of the isolates whereas the largest MGT5 ST (ST900) was assigned to 5.38% (236/4,388) of the isolates.

**Figure 2.**
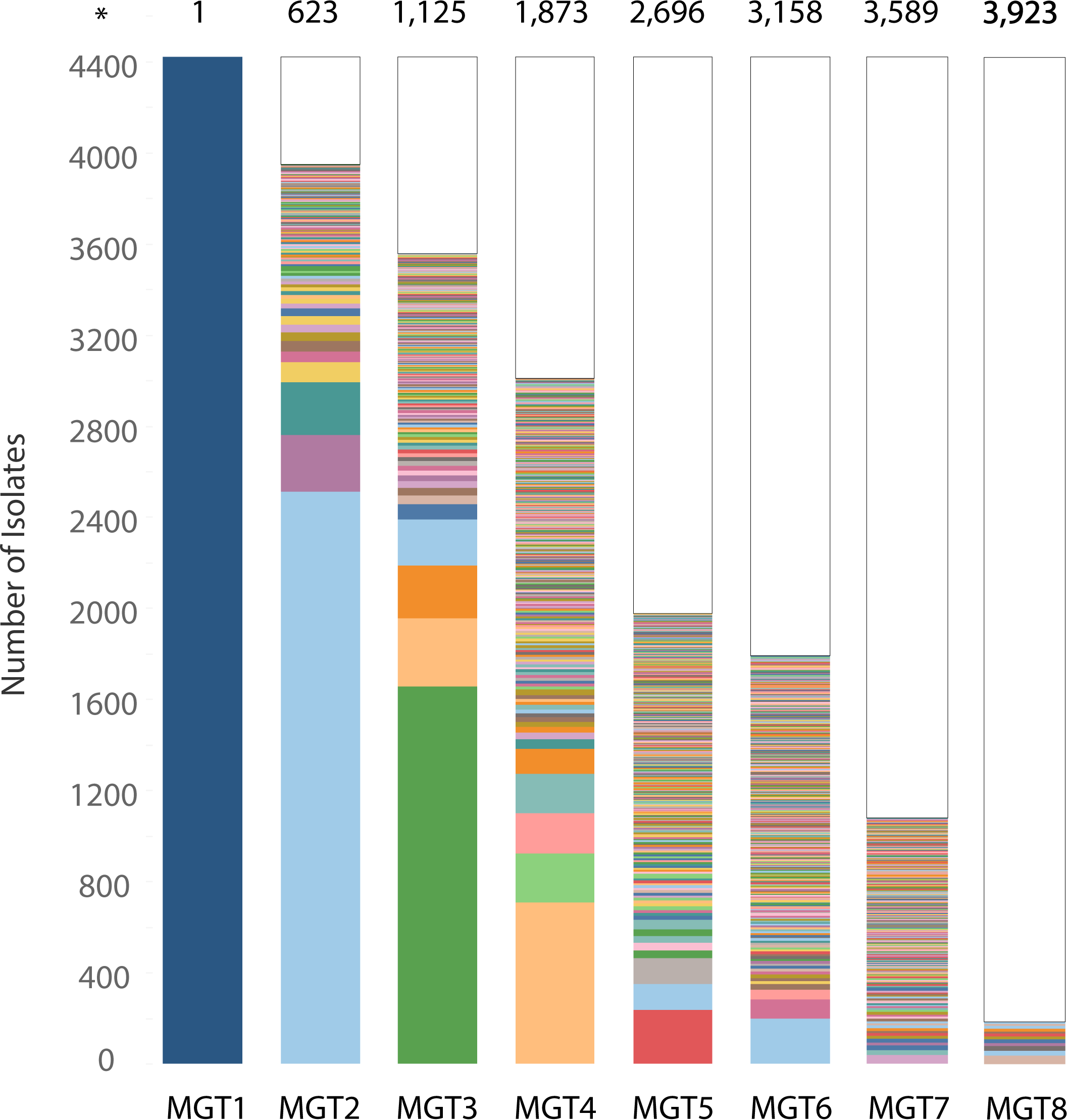
The size distribution of MGT2 - MGT8 STs assigned to MGT1 ST8 isolates. The MGT was used to classify isolates in the MGT1 (MLST) ST8 dataset (n=4,388). The sizes of STs are represented as coloured bars at each MGT level. STs that are assigned to a single isolate (singleton) were collapsed into a single white box. The total number of STs assigned at an MGT level are labelled above (marked by asterisk). Visualised in Prism (v9.3.1)

By MGT2, there were 24 STs with more than 10 isolates. Isolates were sampled from 2008 to 2019, and MGT2 STs varied in frequency over that time (Figure 3), with STs persisting over multiple years and STs sampled only in a single year. MGT2 ST1 was the only MGT2 ST that was sampled in all years and was by far the largest. In eight of those years (2008 - 2011 and 2016 - 2019), MGT2 ST1 was assigned to more than 70% of isolates (Figure 3 shown in dark blue). The second largest type, MGT2 ST199, was sampled only in 2012 and 2013 (Figure 3 shown in light blue). The third largest ST, MGT2 ST114, was similar to MGT2 ST1 and was sampled in all years except 2014 and 2017 (Figure 3 shown in orange). There was a considerable difference between MGT2 ST114 and MGT2 ST1 based on the frequency of isolates over time. MGT2 ST114 had 45.17% (262/580) of isolates sampled in 2015, and an average of 2.25 isolates for the remaining eight years (2008 – 2013, 2016 and 2018).

**Figure 3.**
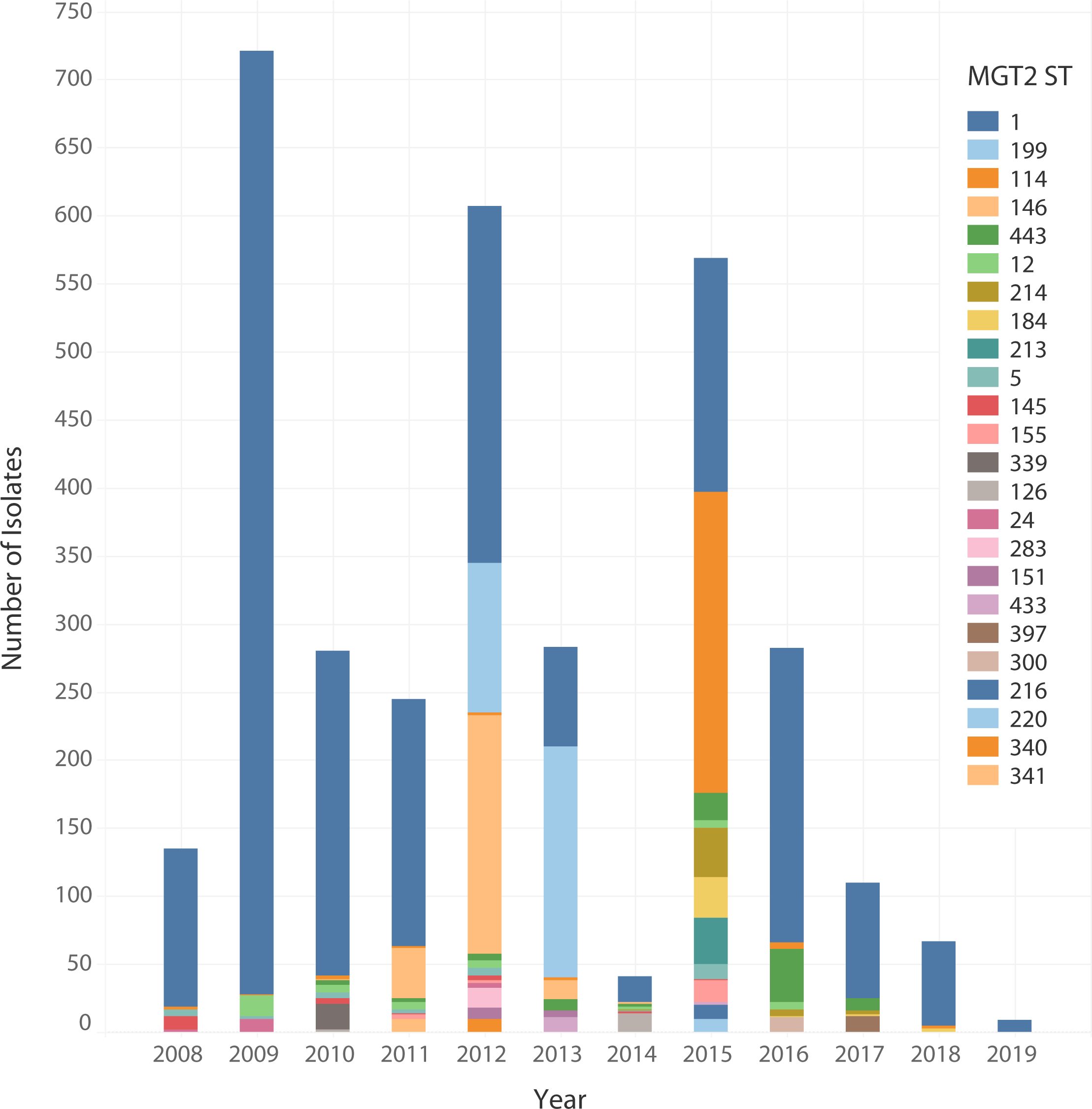
Distribution of MGT1 ST8 isolates by year and MGT2 ST. The temporal distribution of 3,478 MLST ST8 isolates coloured by MGT2 ST. The size of each bar represented the number of isolates assigned to each MGT2 ST. MGT2 STs in the figure legend were organised in descending order of frequency. Visualised in Tableau (v9.1) (49).

MGT typing was compared with *Spa* typing for MGT1 ST8 isolates. The *spa* types of MGT1 ST8 isolates were predicted *in silico*. A *spa* type was predicted for 99.91% (3,475/3,478) of isolates and there were 121 unique *spa* types (Figure 4A). A large number of predicted *spa* types were small in size. Of the 121 *spa* types, 104 were assigned to less than 10 isolates, which cumulatively represented 6.15% (214/3,478) of the MGT1 ST8 dataset. The majority of these 104 *spa* types (54.81%, 57/104) were assigned to a single isolate (Supplementary Table 1) The *spa* types assigned to 10 or more isolates were selected for comparison with MGT2 STs. There were 17 *spa* types selected that together typed 93.85% (3,264/3,478) of MGT1 ST8 isolates (Figure 4B). The majority of isolates were assigned to *spa* type t008 (66.04%, 2,297/2,478). The next two largest *spa* types were t211, t064 which typed 312 and 309 isolates, respectively. The remaining 14 *spa* types were all assigned to less than 100 isolates.

**Figure 4.**
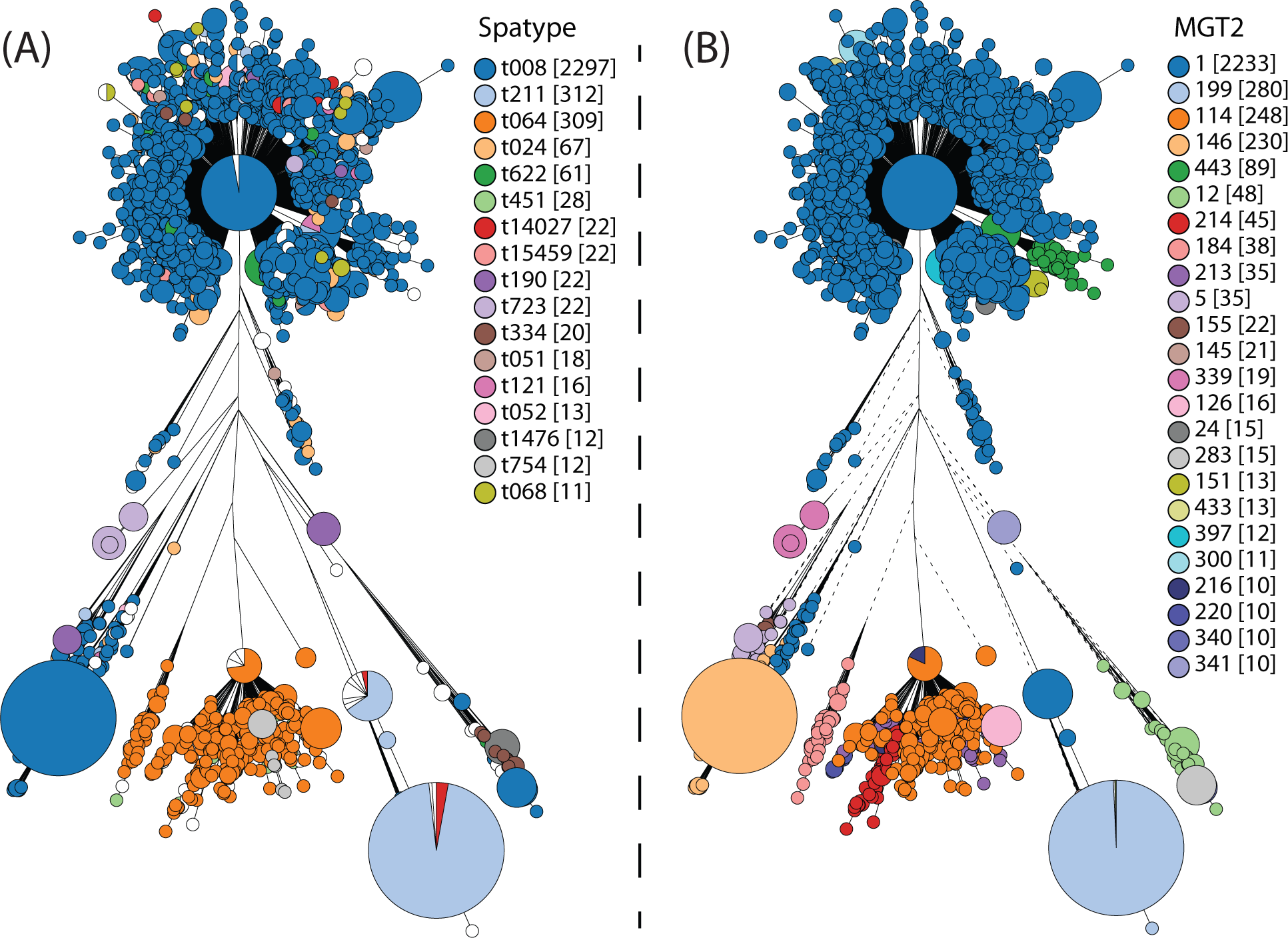
Comparing *Spa* and MGT2 typing within MGT1 ST8. The MGT1 ST8 phylogeny (*n=3,488*) was compared with MGT2 and *Spa* typing. The phylogeny was generated with GrapeTree (v1.0.0) which used the rapid neighbour joining algorithm to process MGT8 (cgMLST) allele profiles (29). Each node was an MGT8 ST and nodes were coloured by *spa* types (A) and MGT2 STs (B). Only *spa* types and MGT2 STs assigned to 10 or more isolates were shown. The frequency of each type was shown in square brackets.

The division of the MGT1 ST8 dataset by MGT2 STs and *spa* types was compared (Figure 4, Supplementary Figure 1). Eight MGT2 STs had a single *spa* type. MGT2 ST283, ST300 and ST340 isolates were *spa* type t008, MGT2 ST126 and ST216 were *spa* type t064, MGT2 ST341 was *spa* type t190, MGT2 ST339 was *spa* type t723, and MGT2 ST397 was *spa* type t622. Other MGT2 STs had a predominant *spa* type with one or more other *spa* types.

### MGT investigation of MGT1 ST8 asymptomatic *S. aureus* colonisation

We used the WGS data from a study that tracked *S. aureus* colonisation of different body sites of the same patients (30) to demonstrate the usefulness of multiple level resolution of MGT. The 82 *S. aureus* MGT1 ST8-USA300 isolates from a cohort of 29 patients were typed by MGT, 24 of which had more than one isolate. A maximum likelihood phylogeny was constructed using core SNPs (Figure 5).

**Figure 5.**
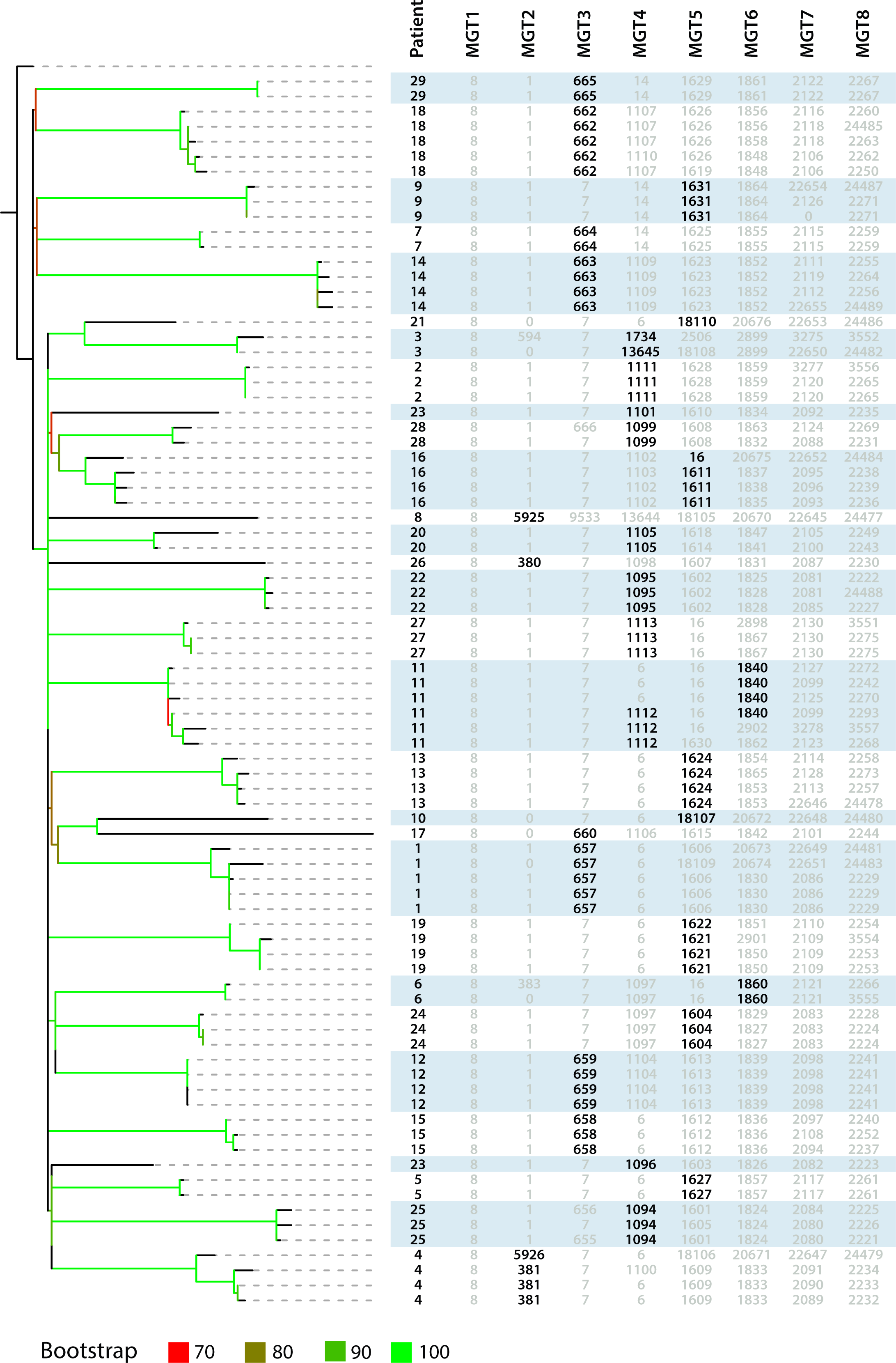
MGT1-MGT8 classification of MGT1 ST8-USA300 isolates from the same patient. A collection of MGT1 ST8-USA300 isolates (n=82) were sampled from 29 patients and classified by the MGT. The MGT1-MGT8 STs and anonymised patient identifier for each isolate were aligned next to the phylogeny. STs that were selected to group isolates of the same patient were bolded. STs in grey were not used to describe multiple isolates of the same patient. The phylogenetic tree was generated using maximum likelihood based on variations in core SNPs. Branches were coloured based on bootstrap support values per colour legend. Branches with bootstraps below 70 were not coloured and remained black. Phylogeny was visualised and annotated with metadata with iTOL (v6) (35).

Using levels MGT2 to MGT6, a total of 34 STs were identified that were specific to one of the 24 patients. In 18 patients (1, 2, 5 - 10, 12 - 15, 17 - 18, 20 – 22 and 24 - 29) all isolates were typed with a specific MGT ST (Figure 5). The remaining 6 patients (3, 4, 11, 16, 19, and 23) each had isolates typed by two specific STs (Figure 5). Only Patient 11 required STs from 2 different MGT levels to describe all isolates with 3 isolates each by MGT4 ST1112 and MGT6 ST1840.

We further identified the number of patient specific STs at each MGT level and what percentage of total isolates were assigned to them. STs from MGT1 to MGT5 were unable to assign all isolates to STs that were specific to one patient (Table 3). MGT6 was the first level to only contain patient specific STs. At MGT6 13 patients required two or three STs to group all isolates. There were seven patients (4, 20, 22 - 24, and 27 - 28) that had each two MGT6 STs, and six patients (1, 11, 13, 16, and 18 - 19) that each had three MGT6 STs which grouped all isolates.

**Table 3.** MGT6 was the lowest level to group all isolates of the same patient.

| Level | Patients Described | Patient Specific STs | Isolates in Patient Specific STs (% of Dataset)# |
| --- | --- | --- | --- |
| MGT1 | 0 | 0 | 0 |
| MGT2 | 5 | 6 | 9.76 |
| MGT3 | 11 | 12 | 36.59 |
| MGT4 | 17 | 21 | 53.66 |
| MGT5 | 27 | 35 | 86.59 |
| MGT6 | 29 | 49 | 100 |
| MGT7 | 29 | 61 | 100 |
| MGT8 | 29 | 69 | 100 |

### MGT application to investigation of MGT1 ST239 transmission in a hospital setting

In 2020, Binary typing was used to characterise the spread of MRSA throughout Concord Hospital (Sydney, NSW, Australia) (31). There were 238 MRSA isolates collected and typed using multiplex PCR-reverse line blot binary typing (referred to as Binary typing) between July 2012 and November 2014. Ninety-seven of the isolates were sequenced and 18 isolates typed as (MGT1) ST239-SCC*mec* III.

The ST239-SCC*mec* III isolates were divided into three BTs: 280841, 280973 and 281997 and typed using MGT (Figure 6). Four MGT STs from MGT2 to MGT5 described the same sets of isolates described by BT (Figure 6). MGT2 ST1294 was equivalent to BT281997 and MGT3 ST2063 was equivalent to BT280973. MGT5 separated BT280973 into MGT5 ST4115 and ST4123 (Figure 6 outlined in grey), representing isolates acquired in different hospital areas. MGT5 ST4123 was acquired in the intensive care unit and general hospital and MGT5 ST4123 was acquired within the burns operating theatre and ward. Within the burns ward, BT281997 (MGT2 ST1294) was divided into MGT8 ST5758, which was patient acquired, and MGT8 ST5743, which was environmental. Thus, MGT STs from MGT8 could distinguish patients isolates from those sampled from the environment (Figure 6 outlined in grey). BT280841 was the only BT unable to be described by a ST from a single MGT level (Figure 6 shown in green). BT280841 was separated into MGT4 ST3013 and MGT5 ST4153,with the latter only contained burns ward environment isolates ( Figure 6 indicated by red and orange boxes). MGT4 ST3013 contained isolates sampled from patients and the environment with the patient isolates distinguished by MGT5 ST4144 (outlined in grey).

**Figure 6.**
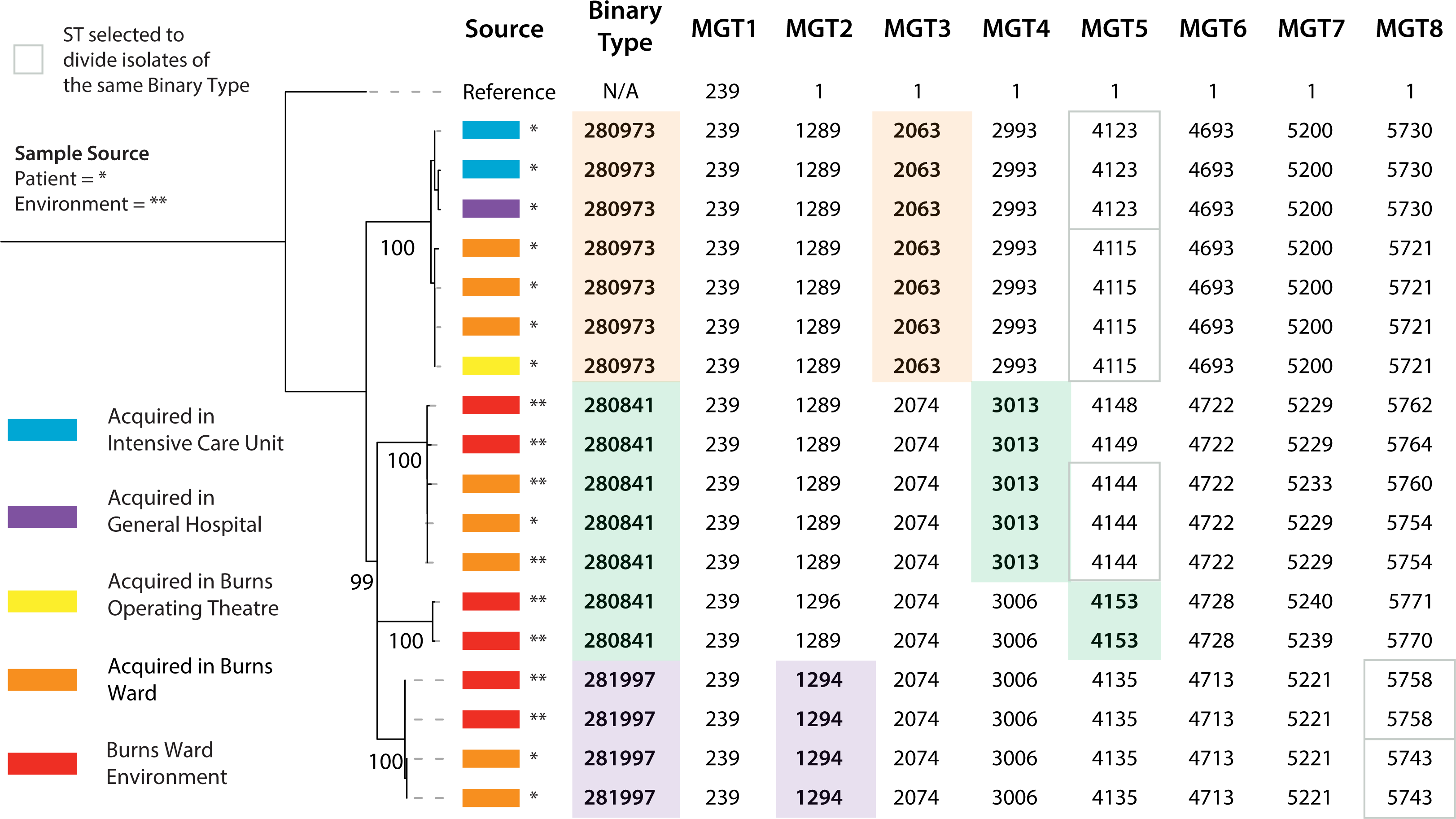
MGT classification of MGT1 ST239-SCC*mec* III spread within Concord Hospital. The spread of MGT1 ST239-SCC*mec* III through Concord Hospital (Sydney, Australia) was described by MGT and Binary typing. Three binary types were coloured as orange, green and purple. The same colouring scheme was used to mark MGT STs that were selected to group isolates the same as binary types. MGT STs that divided binary types in concordance with phylogenetic structure were outlined with a grey box. Patient sampled isolates were marked with an asterisk and isolates sampled from the environment were marked with a double asterisk. Isolates metadata was visualised with the phylogeny using iTOL (v6) (35).

## Discussion

*S. aureus* is an opportunistic pathogen that is the leading cause of skin and soft tissue infections worldwide (1). The carriage of *S. aureus* is strongly associated with the onset of *staphylococcal* infections (39). The development of genomic classification technologies that characterise *S. aureus* population structure and transmission is essential to designing and implementing control and prevention strategies. In this study, an MGT scheme was developed that classified all publicly available *S. aureus* WGS data (*n=50,481*). The MGT was used to investigate both large and small scale *S. aureus* genomic epidemiology using published data as case studies.

### MGT for the standardised genomics-based classification of *S. aureus*

In 2007, the European Society of Clinical Microbiology and Infectious Diseases released guidelines for the development of novel classification technologies (40). These guidelines emphasised the importance of creating new typing technologies that define a standardised nomenclature, offer flexibility in typing resolution and assign types that are interpretable and easy communicable. The *S. aureus* MGT has eight levels filled with loci from the species core genome. The increasing size of each MGT level offers flexibility in typing resolution while maintaining the standardised ST nomenclature at each level. These standardised MGT STs are easily interpretable to encourage the global communication of *S. aureus* large and small scale genomic epidemiology.

The range of typing resolutions make the MGT a useful epidemiological tracing tool. At lower resolution levels, STs tend to be larger, longer lived and more widely distributed across the globe. Additionally, the global distribution of MRSA SCCmec types can also be described. As the resolution increases STs become smaller, persist for less time, and become continent or country specific. At the highest resolution, schemes with equivalent resolution to MGT8 (cgMLST) have been sufficiently precise to uncover chains of transmission and outbreak origins (16, 41, 42). A benefit of MGT classification of closely related isolates is the highest resolution MGT8 can distinguish individual isolates, and the lower resolution levels can place these isolates into a broader epidemiological context.

### MGT described the large-scale population structure of a globally disseminated clone

MGT1 ST8 is a global ST that spread in the Americas and Europe but has also been reported in Africa and Asia (9, 43–45). Using MGT2 we can describe country specific STs for the USA (MGT2 ST114, ST199 and ST443) or the UK (MGT2 ST146). By contrast we can also identify STs that are still globally distributed even at the significantly higher resolution of MGT4 (MGT4 ST6 and ST14). Within MGT1 ST8 the USA300 clone is of particular interest due to its hypervirulence and association with spa type t008 and SCCmec type IV. MGT2 ST1 is the dominant MGT2 ST within MGT1 ST8 and is dominated by SCCmec type IV, spa type t008 isolates. MGT2 STs also describe other large spa types such as t064 (MGT2 ST114), associated with Africa (46) and t211 (MGT2 ST199). MGT provides simpler descriptions of many MGT1 ST8 clades than spa types. Several spa types that are polyphyletic are divided into several smaller but more phylogenetically congruent MGT2 STs such as spa t064 being divided into MGT2 ST114 and ST184. Additionally, there are many small spa types that can be more usefully grouped into larger types using MGT2 STs.

### MGT STs can uniquely describe isolates colonising the same patient within ST8-USA300

The carriage of *S. aureus* is an important risk factor when predicting MRSA onset and 50-80% of MRSA infections are caused by isolates already carried by the host (47). The ability for WGS-based classification to type isolates sampled from the same patient can describe the genomic epidemiology of persistently colonising *S. aureus*. The flexibility in MGT typing resolution allowed STs from multiple levels to describe persistent colonisation of patients. As showcased using MGT1 ST8-USA300 isolates, the STs from MGT2 to MGT6 were able to uniquely group MGT1 ST8-USA300 isolates of most patients by using multiple typing resolutions, the majority of patients (24/29) had one unique MGT ST identifier. Combination of two or more levels would allow description of origin and diversity within a patient.

### MGT STs described MGT1 ST239-SCC*mec* III spread throughout Concord Hospital, Sydney

The MGT was used to describe the spread of MGT1 ST239-SCC*mec* III isolates in a hospital from a previous study (31). The multiple resolutions were able to describe both spread between wards and identify isolates that colonised patients or environments within a ward. The STs from higher MGT levels separated isolates that colonised the hospital environments from the patients admitted to that ward, showing that higher MGT resolutions can separate MGT1 ST239-SCC*mec* isolates independently evolving within a hospital ward. Comparing with binary typing, the flexibility in MGT typing resolution divided binary types into STs that better described *S. aureus* acquisition and transmission. The higher resolutions can distinguish more closely related isolates within a hospital ward such as those colonising a hospital environment and those colonising an admitted patient. The application of the MGT to describe MGT1 ST239-SCC*mec* III spread within a hospital acts as a proof-of-concept investigation and can be applied to any *S. aureus* control within clinical settings.

#### 1.1.1 Limitations of this study and MGT classification of *S. aureus*

Investigating *S. aureus* genomic epidemiology in this study used publicly available *S. aureus* WGS data and metadata. The availability of accurate metadata is essential to interpreting *S. aureus* WGS analysis. Almost all temporal metadata for the investigation of MGT1 ST8 large-scale population structure was sourced from the NCBI BioSample dataset. Interpretation of *S. aureus* WGS data is also complicated when metadata is not available which prevents the detailed description of species wide *S. aureus* genomic epidemiology. The agreement of international standards for epidemiological metadata such as country of origin, year of isolation and host and disease-causing status, would enable better use of MGT to describe the genomic epidemiology of *S. aureus*.

A technical challenge when using the MGT to describe *S. aureus* genomic epidemiology is the definition of singleton STs. The division of *S. aureus* genomes into singletons prevents describing the genetic relationship between isolates. The higher MGT levels that offer higher resolution are expected to increasingly separate isolates into singletons. However, when the lower MGT levels also assign isolates into singletons the MGT is challenged to compare the genetic similarity of isolates at lower and/or higher levels. The MGT2 classification of MGT1 ST8 defined 9.80% (430/4,388) of isolates as singleton STs which were excluded in further investigation. It is possible the isolates in these singleton STs were closely related to the isolates in one of the other large MGT2 STs. The MGT2 level had 21 core loci and an isolate assigned a singleton ST needed to only carry a unique allele for one of those 21 loci. These singleton STs are referred to as single-linkage variants which have previously been shown to arise more frequently in bacterial species with higher SNP mutation rates (such as *S. aureus*) (48). In *S. aureus* MGT online database, clonal complexes were also generated at each MGT level, to enable users to view the clonal complex of singleton STs at each MGT level.

### Conclusion

This study developed a MGT for the flexible, stable and standardised genomic classification of *S. aureus*. The MGT consisted of eight levels providing flexibility in typing resolution. MGT described the large-scale population structure of the species including global, continent specific and country specific STs and their relationship to MRSA lineages and SCCmec subtypes. Within the globally disseminated MGT1 ST8, MGT2 was able to match or improve upon the commonly used spa typing method. Using a combination of higher resolution MGT levels, MGT STs were able to describe isolates from each patient precisely from a study of persistent colonisation. Finally, in a hospital outbreak, MGT STs from lower levels grouped isolates that had spread between hospital wards while higher MGT levels assigned STs that distinguished isolates within the same ward. The MGT was able to describe the genomic epidemiology of *S. aureus* from large, long lived, global types to small short lived STs only found in a single patient. The *S. aureus* MGT publicly available (https://mgtdb.unsw.edu.au/staphylococcus/), updated daily and allows both public and private user submissions. The *S. aureus* MGT can assist in the tracking of existing and new *S. aureus* clones which is essential when designing prevention and control strategies to lower the disease burden of this important pathogen.

## Notes

### Competing Interest Statement

The authors have declared no competing interest.

