## Supplementary Figure 1 for "Characterising *Staphylococcus aureus* genomic epidemiology with Multilevel Genome Typing"

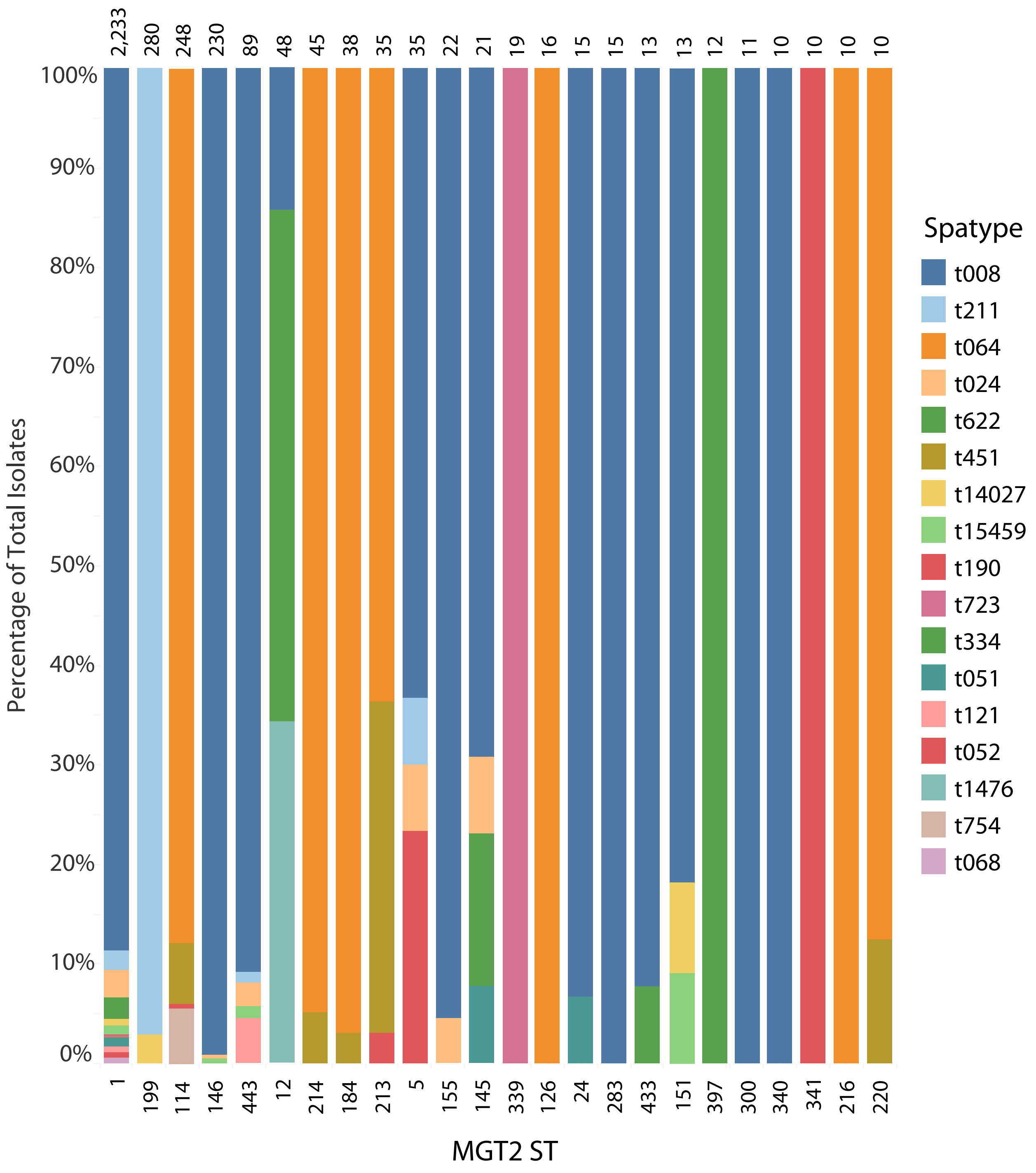


Supplementary figure 1. Comparing MLST ST8 classification between MGT and Spa typing. MLST ST8 isolates were classified with both MGT2 STs and spa types. Isolates were organised into 24 MGT2 STs and then coloured by spa type. The presented spa types were predicted in silico and only spa types assigned to 10 or more isolates were included. For each MGT2 ST, the size of each spa type was the percentage of total isolates. The total number of isolates was labelled above each MGT2 ST. Columns were organised by total number of isolates in descending order. MGT2 STs and spa types were visualised in Tableau (v9.1).
