## Supplementary Methods for "Characterising *Staphylococcus aureus* genomic epidemiology with Multilevel Genome Typing"

**1.1 Calculating the size of each level**

The number of loci per MGT2-7 level was calculated using a whole genome SNP mutation rate and selecting the amount of DNA expected to generate a single SNP in fixed timeframes (**Supplementary Dataset 1, Supplementary Methods Table 1**) (284, 366, 382, 464).

**Supplementary Methods Table 1.** Target time periods for each MGT level and the corresponding basepair length needed to achieve one mutation in that time period.

| **Level** | **Level Time Period (Years)** | **Target Size of Level (basepairs)** |
| --- | --- | --- |
| MGT2 | 20 | 17,559 |
| MGT3 | 10 | 35,118 |
| MGT4 | 5 | 70,236 |
| MGT5 | 2 | 175,589 |
| MGT6 | 1 | 351,178 |
| MGT7 | 0.5 | 702,356 |

**1.2 Calculating locus filtering metrics**

The following metrics were then calculated for each locus of the core genome: dN/dS ratio, locus quality, homopolymers, tandem repeats and phage coding regions (**Supplementary Datasets 2 to 6.**). Calculated metrics unique to this study were a rate of allelic change per kilobase and a predicted number of recombination events per locus (**Supplementary Datasets 7 and 8, Supplementary Scripts 1 and 2**).

**1.3 Calculating the rate of allelic change per kilobase**

The rate of allelic change for a core locus was used as a proxy for measuring the rate of evolution. The rate represented the number of alleles per kilobase for a locus. Per core locus, the rate of allelic change was the average of two test datasets each with 10,000 genomes. Each dataset represented the seven-gene MLST diversity of species. An in-house python script sampled each ST proportionally based on that STs frequency in the species dataset (**Supplementary Script 1**). A single isolate was selected from all singleton STs. Allele profiles for the test datasets were generated using the MGTdb Allele Calling pipeline (150). Prism was used to compare the distribution of alleles per core gene (313). The means of test-dataset one and test- dataset two were compared with a Man-Whitney U test. Significance between means was interpreted with an alpha threshold of 0.95. The number of alleles per core gene was averaged between the two datasets and normalised per kilobase.

Core loci were separated into percentiles based on the distribution of rate of allelic change per kilobase. The distribution of values was tested for normality using a Shapiro-Wilk test. A p- value less than or greater 0.05 and alpha value higher than 20 indicate a distribution departing from normality.Genes were categorised into the following percentiles within a negative binomial distribution: 10th, 20th, 30th, 40th, 50th, 60th, 70th, 80th, 90th and 100th.

**1.4 Calculating the predicted number of recombination events per locus**

The number of recombination events impacting each of the core loci was predicted. A core SNP alignment was generated for representative dataset using Snippy (126). Isolates of the representative dataset were aligned to *S. aureus* COL. The regions predicted to be under the influence of recombination were identified with RecDetect (310). A strict prediction model for high recombination species was selected. For each core locus, the number of recombinant regions that overlapped was counted with an in-house python script (**Supplementary Script 2**). Both partially and completely overlapping regions were identified. A partial overlap was a recombinant region that overlapped the start or end positions of a locus. A complete overlap was a recombinant region within the start and end positions of a locus. The number of recombination events per core locus was visualised in Prism (313).

**1.5 Separating the core loci into preferences**

All calculated metrics were the basis for separating the core loci with a preference-based system (148, 149, 347). An in-house python script separated the core loci into preferences 1- 12 using the filtering thresholds in **Supplementary Methods Table 2 (Supplementary Script 3)**. MGT2-7 were filled by randomly selecting loci from the lowest preference. The levels were filled in ascending order with loci that were separated by the distances shown in **Supplementary Methods Table 3**. The loci from the next highest preference were selected when an MGT level required additional loci to meet the scheme size **(Supplementary Script 3)**.

**Supplementary Methods Table 2.** Preference classifications assigned to each locus depending on their characteristics

| **Preference** | **Allelic Rate Percentile** | **dN/dS Ratio Percentile** | **Missing (%)** | **Problematic Allele Calling (%)** | **Recombination Events** | **Homopolymer** | **Tandem Repeats** | **Phage Region** |
| --- | --- | --- | --- | --- | --- | --- | --- | --- |
| 1 | 10 | 30 | <=1 | <=1 | <=15 | False | False | False |
| 2 | 20 |  |  |  |  |  |  |  |
| 3 | 30 |  |  |  |  |  |  |  |
| 4 | 40 | 50 |  |  | <=20 |  |  |  |
| 5 | 50 |  |  |  |  |  |  |  |
| 6 | 60 | 90 |  |  |  |  |  |  |
| 7 |  |  |  |  |  |  |  |  |
| 8 | 70 |  |  |  | <=25 |  |  |  |
| 9 | 90 |  |  |  |  |  |  |  |
| 10 | 95 | 100 |  |  |  |  |  |  |
| 11 | 100 |  |  |  |  |  |  |  |
| 12 |  |  |  |  |  | True | True | True |

Note: The core loci were divided into preferences 1-12. Multiple metrics for each core locus were considered when assigning a preference. The filtering thresholds used to rank loci into preferences were shown.

**Supplementary Methods Table 3.** Minimum allowable distance between isolates.

| **MGT level** | **Separation Between Loci (bases)** |
| --- | --- |
| MGT2 | 20,000 |
| MGT3 | 12,500 |
| MGT4 | 5,000 |
| MGT5 | 3,500 |
| MGT6 | 500 |
| MGT7 | 0 |
